# Voice and linguistic features support early attentional filtering of irrelevant stimuli

**DOI:** 10.64898/2026.09.17.752368

**Authors:** Sahil Luthra, Eric Parker, Marysia Brown, Gidey Gezae, Hee So Kim, Wusheng Liang, Abigail Noyce, Barbara G. Shinn-Cunningham

## Abstract

In cocktail party-type environments, listeners must segregate simultaneous acoustic sources and select one for processing. Behavioral studies have established that both low-level voice differences (e.g., pitch) and high-level linguistic differences (e.g., word content) aid these processes. Electroencephalography (EEG) studies have demonstrated that the brain filters voices based on low-level features early in sensory processing, but it is unclear whether linguistic differences also support early filtering. In this EEG study, listeners attended to a lateralized auditory stream of target syllables while ignoring distractor stimuli from the opposite hemifield. In contrast to previous auditory attention studies, target stimuli were always attended and statistically identical across conditions; instead, we manipulated the distractors. Specifically, we manipulated whether targets and distractors were spoken in the same voice or different voices, as well as whether they were composed of the same or different linguistic content. Critically, distractors were temporally offset from targets, enabling an investigation of whether acoustically matched, always attended target stimuli are differentially encoded as a function of surrounding context. Differences in either voice or linguistic features supported target recall: Recall was poor only when both streams were in the same voice and had similar linguistic content. However, voice and linguistic features had independent, additive effects on neural processing. Specifically, target dissimilarity in acoustics and linguistic content both contributed to more robust, larger-magnitude target-evoked EEG responses. Overall, results demonstrate that both linguistic dissimilarity and low-level acoustic dissimilarity improve early attentional filtering of distractors in multi-source acoustic scenes.

**Graphical Abstract:** 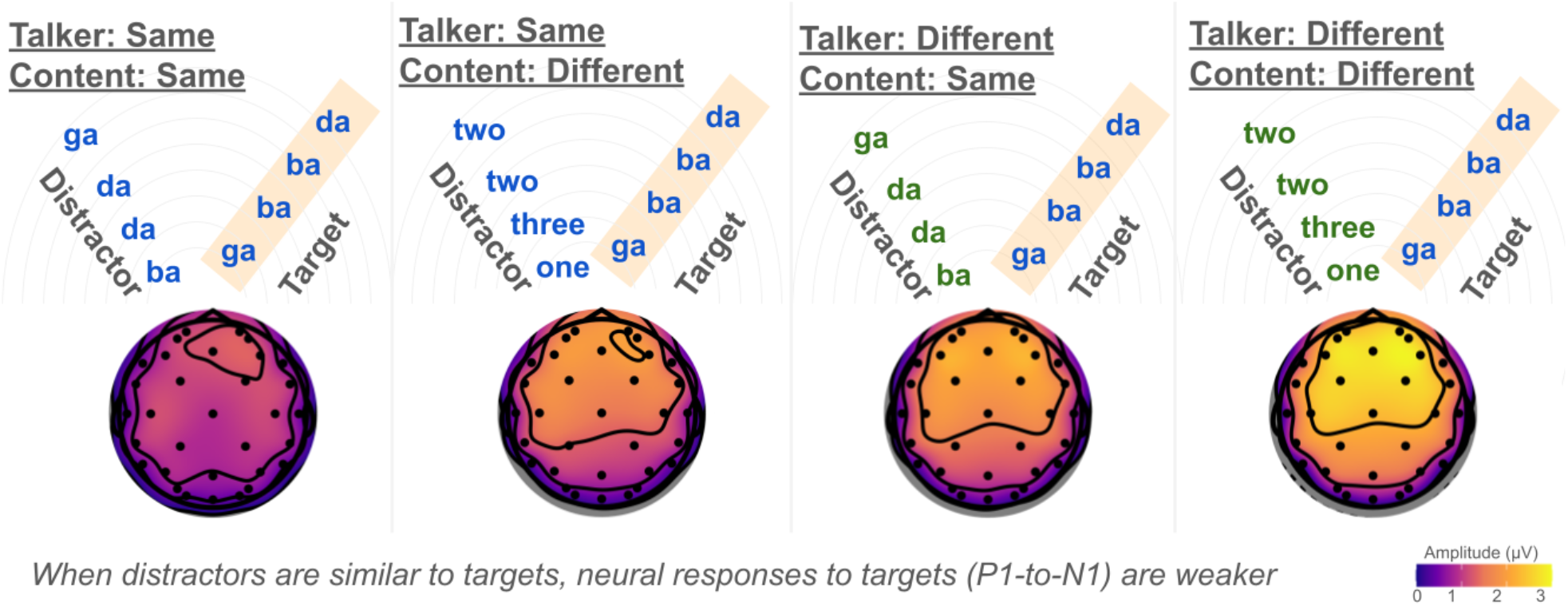

**Highlights:**

- We varied the voice and semantic content of distractors in a cocktail party task
- Auditory targets were statistically identical and always attended across conditions
- Target recall suffered only when streams were similar in both voice and content
- But voice and linguistic features independently shaped attentional EEG responses
- Similarity in either voice or content weakened target-evoked neural responses

## 1. Introduction

To efficiently process the barrage of sensory information we often encounter, top-down attention filters out irrelevant information, prioritizing what is most important (Desimone & Duncan, 1995). However, we lack an understanding of whether the neural mechanisms that support attentional filtering differ depending on which specific features support attention, including low-level features such as color (in vision) or pitch (in audition) and higher-level representations of meaning.

Consider selectively attending to a single conversation in a cocktail party environment (Cherry, 1953). Listeners can filter out distracting conversations based on various features, including differences in location, pitch, timbre, and linguistic content. Electroencephalography (EEG) studies show that filtering out distractors using pitch influences the early encoding of target stimuli, such as the P1, N1, and P2 components of the auditory event-related response (ERP; e.g., Snyder et al., 2006), but it is unclear whether semantic or linguistic differences between streams similarly support early filtering.

Voice and linguistic features may support attentional filtering through distinct mechanisms. Voice cues contribute to *segregating* the input into distinct auditory objects as well as *selecting* a single source for attentional prioritization (Shinn-Cunningham et al., 2017). By contrast, access to linguistic cues requires that an utterance have already been processed, at least partially, by relatively high-level cortical areas; linguistic cues thus likely contribute more strongly to selection than segregation. Accounts of speech perception also differentiate between talker and linguistic attributes of the signal (e.g., Peterson & Barney, 1952; Ladefoged & Broadbent, 1957; Pisoni, 1997; Smith et al., 2002), and these cues are processed by distinct but overlapping neural systems (Hickok & Poeppel, 2007; Luthra, 2021; Maguinness et al., 2018). However, extensive feedback circuitry throughout the brain may allow both features to shape early sensory responses (e.g., Yantis, 2008) via top-down connections from high-level regions.

In this EEG study, we investigated the behavioral and neural mechanisms that allow listeners to filter out distracting auditory streams using voice or linguistic features. Thirty normal-hearing adult listeners were asked to report the contents of one of two competing speech streams (spatialized ±30° left or right) while ignoring the other. Most attentional studies compare neural responses evoked by the same signal when it is attended versus ignored. Instead, in this study, we contrast responses to the same acoustic target stream while varying characteristics of the competing distractor. Specifically, listeners always attended to a target stream comprising four syllables, sampled with replacement from the set {*ba, da, ga*}, spoken by a single voice and lateralized to the same hemifield for each given participant. In a 2×2 factorial design (Figure 1), the distractor stream was either spoken in a different voice or the same voice, and comprised either digits sampled from the set {*1, 2, 3*} (different linguistic content than the target, facilitating selection) or the same syllables as the target (challenging selection).

**Figure 1.**
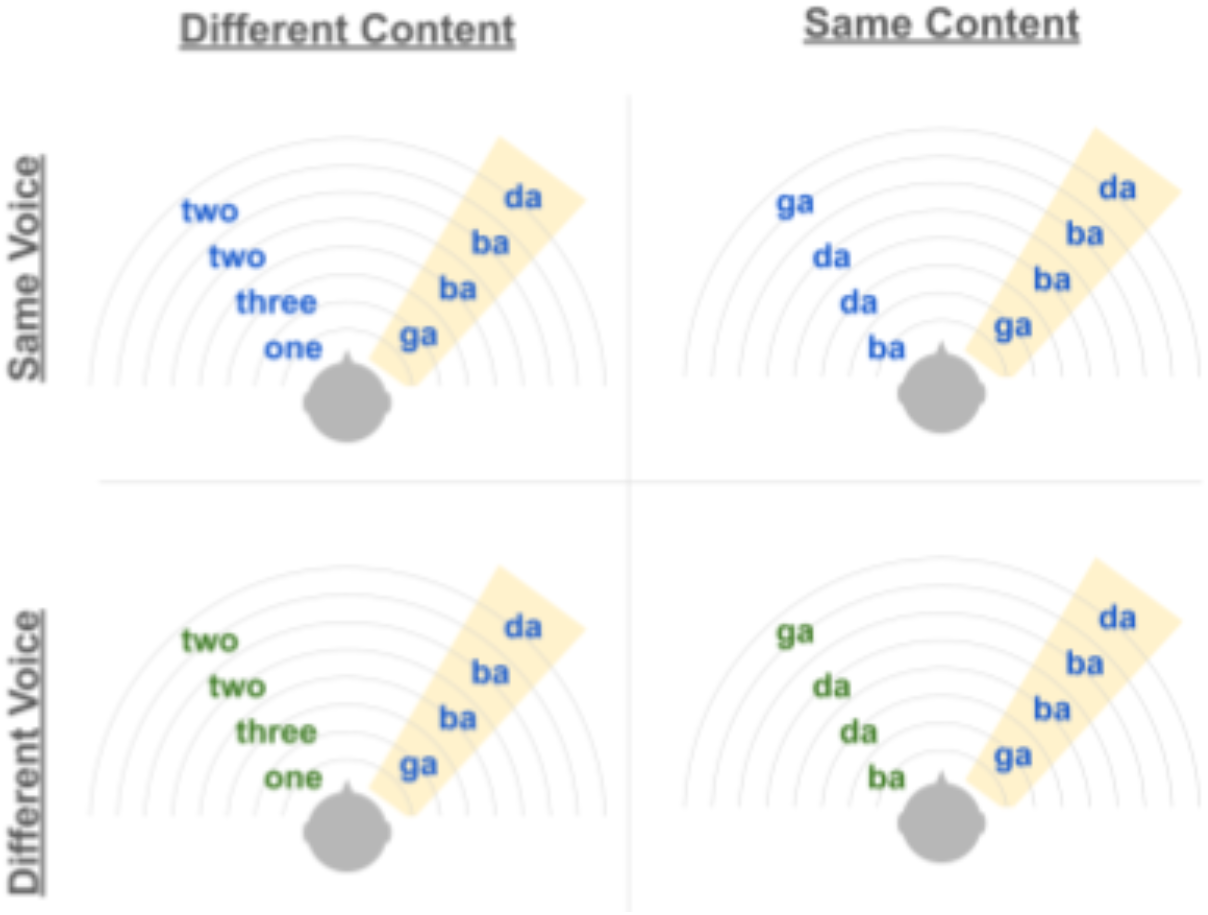
Experimental design. Target syllables (*ba. da. ga* spoken by a male voice in a fixed location) are interleaved with spatially separated distractor stimuli (streams spatialized ±30° azimuth; here, target is on the right). Across conditions, we manipulate whether the distractor stimuli are similar in linguistic content (making target selection difficult) or different, as well as whether distractor stimuli are spoken in the same voice.

Distractors and target streams were temporally interleaved, allowing us to measure behavioral and neural responses to targets that were matched acoustically across conditions and uncontaminated by simultaneous distractor stimuli. By manipulating the context surrounding the statistically identical, always-attended target stream, we assessed how attentional filtering varies with the voice and linguistic properties of a competing stream. We measured how target syllable recall and event-related EEG responses to target stimuli varied based on voice and linguistic similarity between the target and distractor.

We hypothesized that both voice or linguistic differences would aid attentional filtering and improve target recall; we expected recall to be worse when targets and distractors were similar in both voice and linguistic content. We expected that early ERP components would be influenced by voice properties of the distractor, based on prior findings (e.g., Snyder et al., 2006). By contrast, we hypothesized that linguistic differences would not modulate early auditory encoding, given that such differences operate at a higher, more abstract level of auditory processing.

## 2. Materials and Methods

### 2.1. Stimuli

Stimuli (syllables *ba, da*, and *ga*, and the spoken digits *1, 2*, and *3*) were recorded by two native speakers of American English (one female, one male). The duration of each stimulus was set to 300 ms using the Time-Domain Pitch-Synchronous Overlap-and-Add (TD-PSOLA) algorithm (Moulines & Charpentier, 1990), and all stimuli were scaled to a sound level of 70 dB SPL; these preprocessing steps were implemented in in Praat (Boersma & Weenik, 2017; RRID:SCR_016564).

To create trials with competing streams originating from opposite hemifields, each stimulus was spatialized in MATLAB (The Mathworks Inc., Natick, MA, USA; RRID:SCR_001622) using head-related transfer function (HRTF) measurements from a KEMAR dummy-head microphone (Gardner & Martin, 1995). This was achieved by convolving each recording with the left- and right-ear filters corresponding to sources at ±30° azimuth. In pilot testing, ±30° spatial separation was deemed to provide a sufficient cue to enable selective attention in the absence of voice or linguistic differences (i.e., when both target and distractor streams were composed of syllables produced in a male voice), while avoiding overly salient spatial cues that could dominate performance.

Temporally interleaved stimulus sequences were generated in MATLAB. Each trial consisted of two concurrent streams: a target stream and a distractor stream. The target stream always comprised male syllables, whereas the distractor stream varied along two factors: Content (syllables vs. digits) and Voice (same vs. different relative to the target), resulting in a 2 × 2 design (same/different voice × same/different content). Each stream was composed of a sequence of four items drawn randomly, with replacement, from a set of three possible items ({*ba,ga, da*} or {*1, 2, 3*}). Sequences were randomized with the constraint that no sequence contained four identical items (e.g., “ba ba ba ba” or “1 1 1 1”). Target and distractor sequences were generated independently, with the additional constraint that the two streams did not contain identical sequences on a given trial (e.g., “ba ga da da” paired with “ba ga da da”). The two streams were then interleaved to create sequences of eight items such that items alternated between streams (e.g., target-distractor-target-distractor-…), with each item presented for 300 ms and no temporal overlap between streams, resulting in a continuous alternating sequence. On each trial, the target stream either led (first item) or lagged (second item), with equal random likelihood; this factor was counterbalanced within participants. The target stream was presented in either the left or right hemifield (±30° azimuth), with the distractor stream presented to the opposite side; target location was fixed for each participant (i.e., half of the subjects always heard the target from the left and half always heard it from the right).

### 2.2. Participants

Young adults (18-35 years old) were recruited from the Carnegie Mellon University community and were compensated for their time in either cash or partial course credit. All were self-reported native speakers of English with no diagnosed hearing impairments, normal or corrected-to-normal vision, and no known neurological disorders. Participants provided informed consent following the procedures approved by the Institutional Review Board of Carnegie Mellon University. Normal hearing was verified via audiometric screening in a sound-attenuated booth; participants were only eligible to continue to the experimental task if they had pure-tone thresholds at or below 20 dB HL for octave frequencies between 125 and 8000 Hz as well as at 6000 Hz. We recruited 45 participants, of whom 7 failed the hearing screening, 3 did not complete the experiment, and 5 were dismissed due to technical issues. This resulted in a final sample of 30 participants (19 female, 9 male, 2 not reported; mean age: 21, age range: 18-28). This sample size was selected based on a power analysis of pilot behavioral data, which indicated that a sample size of 20 would be sufficient to exceed 80% power for fixed effects of Talker, Content, and their interaction. We opted for a slightly larger sample size based on previous EEG studies from our group.

### 2.3. Procedure

Raw EEG data were collected through a BioSemi ActiveTwo 32-channel EEG system (10/20 layout) using Ag-AgCl electrodes. Common Mode Sensor (CMS) and Driven Right Leg (DRL) electrodes were placed on opposite sides of Cz, and external flat-head reference electrodes were placed on the mastoids. Impedances were maintained at or below 20 mV. Four-bit condition-specific event markers (aligned to audio onset using additional channels in each sound file) were recorded in ActiView simultaneously with EEG data.

The experimental task was implemented using custom code in PsychoPy (Peirce, 2007; RRID:SCR_006571). Audio stimuli were delivered using insert earphones with ER-2 (Etymotic) tips. Trial onsets were marked using a Triggy (Cortech) system. Each participant completed between 14 and 20 blocks of 40 trials each and were given short breaks after each block.

### 2.4. Behavioral Analysis

By-subject by-syllable accuracy data were submitted to linear-mixed effects regression analyses implemented in R (R Core Team, 2025; RRID:SCR_001905) using the *lmer* function in the “lme4” package (Bates et al., 2015). For all analyses, we included random intercepts for each participant and specified a binomial distribution for the data.

The primary analysis tested for fixed effects of Voice (same/different, deviation-coded with a [0.5, -0.5] contrast), Content (same/different, deviation-coded with a [0.5, -0.5] contrast), and their interaction. Follow-up analyses were performed by subsetting the data based on distractor content and fitting reduced models that tested only for an effect of Voice.

We also conducted a series of more complex analyses that tested for interactions with Syllable Position and Target Position (whether the target led or lagged behind the distractor on that trial, see Supplemental Information, Table S1). Similar results were obtained with this more complex analysis; therefore, for simplicity, we collapse across these factors in the main text.

### 2.5. EEG Preprocessing

Preprocessing of EEG data was implemented in EEGLAB (Delorme & Makeig, 2004; RRID:SCR_007292) using custom MATLAB code. Data were referenced to the average of the mastoid electrodes and downsampled to 500 Hz. A zero-phase, first-order Butterworth bandpass filter (0.1 - 50 Hz) was applied to minimize slow drift and myogenic artifact; a notch filter was also applied at 60 Hz to remove line noise. Data were epoched at trial onset by condition, and epoched data were then decomposed via automated ICA labeling (Makeig et al., 1995) and inspected visually; components corresponding to eye blinks and muscle artifacts were removed. Data were re-epoched to generate by-stimulus ERPs (i.e., separate ERPs for each target and distractor item). Preprocessed, epoched data were exported to R for visualization (using the “ggplot2” library; Wickham, 2016) and subsequent analyses.

### 2.6. ERP analyses

ERPs were estimated by averaging the responses to the second and subsequent stimuli on each trial (separately for targets and distractors). The first stimulus on each trial (i.e. the first target on target-leading trials and the first distractor on target-lagging trials) was excluded from this analysis, since (1) trial-initial stimuli are associated with disproportionately large responses and (2) it would not make sense to consider effects of distractor type for the first stimulus on a target-leading trial, as the participant would not yet have experienced any distractors on that trial. We limited our analyses to a set of frontocentral electrodes (FC1, FC2, Fz, Cz) where early auditory ERPs are modulated by attention (e.g., Fleming et al., 2020; Snyder et al., 2006). We limited our analysis of ERPs elicited by target stimuli to those that were correctly identified by participants, but obtained nearly identical results in an analysis that included all target stimuli. Analyses of the time course of ERP responses (for both target and distractor stimuli) are reported in Supplemental Information.

A priori, we were principally interested in attentional effects on early auditory event-related potentials — in particular, the P1, N1 and P2 — elicited by target stimuli. However, because across-stream ISIs were 0, late ERP responses to the previous stimulus token likely overlapped with these early components. To reduce such carry-over effects, our primary ERP measures were peak-to-peak changes in the P1-N1 complex and the N1-P2 complex. In visualizing the results, we opted to baseline correct the ERPs using the average voltage at time zero rather than over a prestimulus window of any duration, as even a short baseline might be skewed by responses evoked by the preceding syllable. P1-N1 and N1-P2 values were only computed for target stimuli, where we observed similar timing of peaks across conditions. For each participant, we defined P1, N1, and P2 peaks using that participant’s target-elicited ERP averaged across all conditions. We first identified the N1 peak using the most negative value between 75 and 190 ms. We then identified their P1 as the most positive-going value that occurred after 30 ms and at least 15 ms prior to the N1, and their P2 as the most positive-going value that occurred at least 15 ms after the N1 and prior to 250 ms.

P1-to-N1 and N1-to-P2 peak-to-peak values were computed for each condition separately, using the average voltage in a 40 ms window centered on each individual subject’s P1, N1, and P2 times as identified above. These were submitted to an ANOVA with factors of Distractor Voice (same/different) and Distractor Content (same/different) implemented using the *ezANOVA* function in the “ez” library (Lawrence, 2016).

## 3. Results

### 3.1. Voice and linguistic differences improve target recall

Overall, participants exhibited strong performance on the target recall task (mean accuracy: 82.4%, SE: 1.6%; chance: 33%; Figure 2). A mixed-effects regression analysis identified a significant effect of Voice (b = -0.143, SE = 0.019, z = -7.543, p < 0.001), a significant effect of Content (b = -0.551, SE = 0.019, z = -26.066, p < 0.001), and a significant interaction between them (b = -0.539, SE = 0.038, z = -14.244, p < 0.001). Follow-up analyses indicated that when selection was easy (the target stream comprised syllables but the distractor stream consisted of digits), there was no additional performance benefit to presenting different voices in the two streams (and in fact, hearing the *same* voice in both streams led to a slight performance advantage in this case, b = 0.127, SE = 0.029, z = 4.416, p < 0.001; accuracy: 86.9% [same voice], 85.6% [different voice]). However, when selection was difficult (both target and distractor streams comprised syllables), hearing the same voice in both streams incurred a significant performance cost (b = -0.414, SE = 0.025, z = -16.76, p < 0.001; accuracy: 75.4% [same voice], 81.8% [different voice]). Overall, results indicate that voice differences aid target recall when competing streams are linguistically similar, but linguistic differences between streams yield strong performance regardless of low-level acoustic similarity.

**Figure 2.**
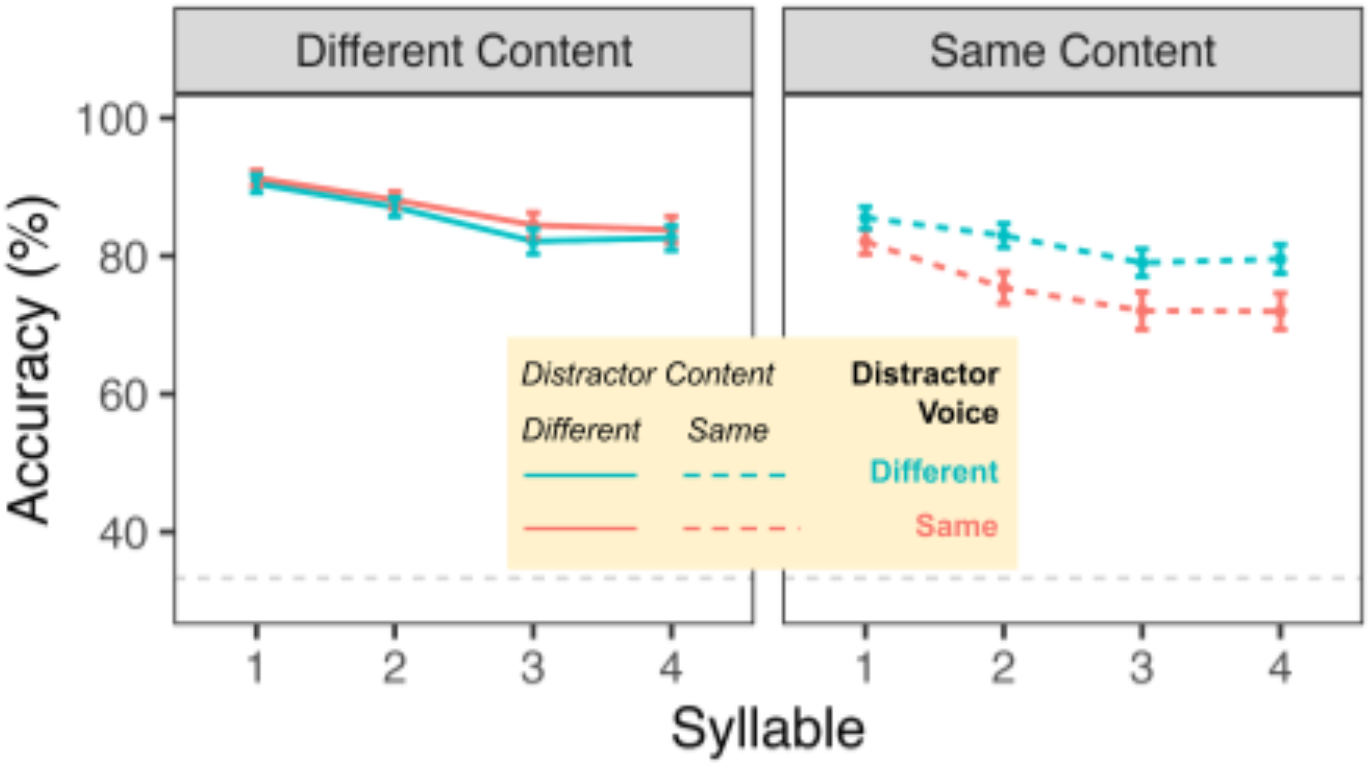
Behavioral accuracy for target syllable recall. When the distractor stream is linguistically distinct from the target (and selection is easy), accuracy of syllable recall is not affected by the voice producing the distractor. However, accuracy is impaired when the distractor stream is similar both in linguistic and voice information. Error bars display standard error. Dashed gray line indicates chance-level accuracy.

### 3.2. Neural encoding of targets is independently influenced by voice and linguistic features

Event-related responses to correctly identified target stimuli are plotted in Figure 3A (green panel, left). Although stimuli were acoustically identical and always attended across conditions, target ERPs were modulated by both the voice and linguistic properties of the competing distractor stream, and these effects were additive. Statistical analyses confirmed that the peak-to-peak differences in the P1-N1 response elicited by target stimuli (Figure 3B) were larger when the distractor stimuli were produced in a different voice (F(1,29) = 42.44, p < 0.001; mean difference: 0.893 μV) as well as when the distractor stream comprised linguistically distinct stimuli compared to the target (F(1,29) = 37.51, p < 0.001; mean difference: 0.734 μV); however, these two factors did not interact (F(1,29) = 0.014, p = 0.907). In contrast to the earlier P1-N1 responses, the peak-to-peak differences in the N1-P2 responses showed no statistical effect of either distractor voice or content (all p > 0.09). The scalp topography of the P1-to-N1 effect (i.e., the change in amplitude from the P1 window to the N1 window) for each condition is shown in Figure 3C; these maps highlight the large peak-to-peak change when the target and distractor differ by two features, the intermediate change when they differ by one feature, and the small change when they differ by one feature.

**Figure 3.**
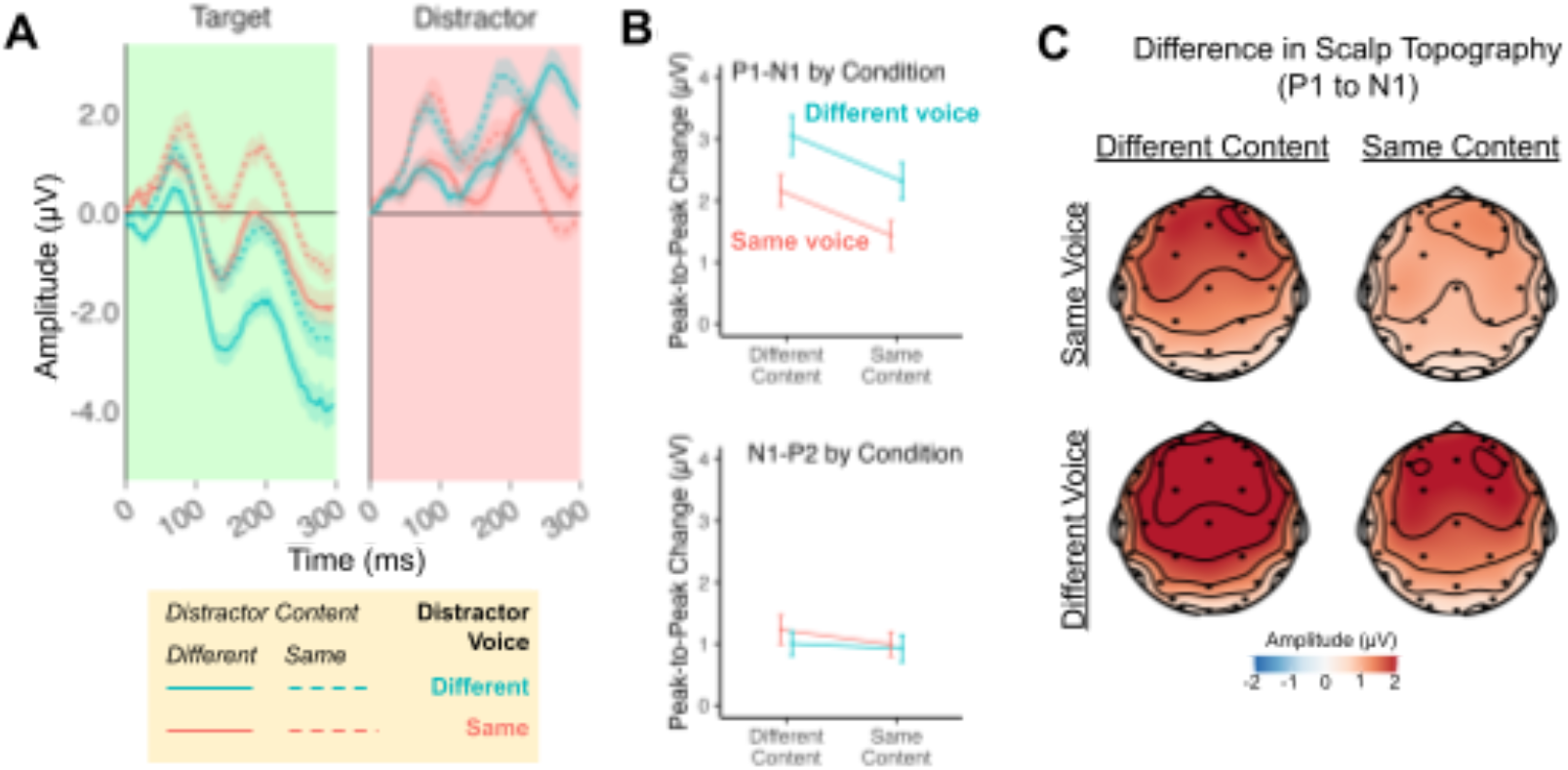
EEG Results. **(A)** Event-related potentials at frontocentral electrodes (averaging across stimulus positions, excluding the first stimulus on each trial) for correctly identified targets (green panel, left) and distractor stimuli (red panel, right). Neural responses to targets are largest when the distractor is maximally dissimilar (solid teal line) and smallest when the distractor matches in both voice and linguistic content (dashed salmon line). **(B)** Peak-to-peak amplitude differences in responses to target stimuli, illustrating that the effects of distractor voice and content influence the P1-to-N1 peak-to-peak response at frontocentral electrodes. **(C)** Maps showing the difference between the P1 scalp topography and the N1 scalp topography for responses to target stimuli in each condition.

## 4. Discussion

We assessed listeners’ ability to selectively attend to a stream of target syllables while ignoring a spatially separated, temporally interleaved distractor stream. Across conditions, we manipulated whether the target and distractor were similar or different in terms of voice and linguistic content. We found that both voice and linguistic differences aided selective attention behaviorally, as these factors interacted to influence recall of target syllables. However, EEG data indicated that voice and linguistic properties of the distractor stimuli had independent, additive effects on the early encoding of target stimuli, modulating the peak-to-peak amplitude between the P1 and N1 components.

Traditionally, studies of attention manipulate the focus of attention, examining responses to stimuli when they are attended compared to when they are ignored. In contrast, the present study held constant both the acoustic properties of target stimuli and the attentional focus across conditions, and instead manipulated the properties of surrounding distractor stimuli. Crucially, target and distractor stimuli were offset both temporally and spatially, allowing us to measure ERPs to target stimuli with minimal energetic masking from distractors. By varying the context in which target stimuli were encountered, we were able to probe the mechanisms that support listeners’ ability to tune out irrelevant streams. We found that early neural responses to target stimuli were modulated by the nature of surrounding distractors. We suggest that the P1-to-N1 response in this paradigm reflects the attentional filter: When a target is highly similar to a distractor, early filtering is less effective, leading to greater neural interference by the distractor and a weaker target-evoked P1-N1 response.

We hypothesized that distinct mechanisms would support attentional filtering based on voice versus linguistic features, in large part because of how voice and linguistic cues contribute to auditory segregation versus auditory selection. While past work has suggested that voice cues contribute strongly to segregation (Assmann, 1999; Brungart, 2001; de Cheveigné et al., 1997; Shackleton & Meddis, 1992), the impact of voice differences has primarily been examined in studies where linguistic differences between streams are minimized (e.g., both streams consist of stimuli from a shared closed set of words). In this case, even if listeners can successfully segregate the acoustic mixture, it may be difficult to correctly *select* one of two linguistically similar streams for prioritization. Our results suggest that the influence of voice features on auditory attention is modulated by linguistic differences between speech streams and concomitant demands on target selection.

Notably, we observe a striking dissociation between neural encoding and behavior: While EEG data indicate that both voice and linguistic cues support the early filtering of target stimuli, behavioral results indicate that when there are linguistic differences between streams, voice differences do not further enhance recall of target stimuli. That is, a single feature difference between streams (whether linguistic content or voice pitch) is sufficient to guide selective attention. However, because recall accuracy only indexes the outcome of selective attention, our data cannot indicate whether selective attention is more challenging when only one feature differentiates competing streams compared to two features. Future research leveraging pupillometry could provide useful insights as to whether the cognitive effort associated with selective attention depends on the number of features that differ between speech streams.

More generally, the current results provide insight into the neural mechanisms that support top-down auditory attention. The behavioral prioritization of task-relevant stimuli is reflected in neural activity across large swaths of cortex, with responses to attended stimuli generally magnified relative to unattended stimuli (e.g., Liu et al., 2003, 2007; Mesgarani & Chang, 2012; O’Craven et al., 1999; Puschmann et al., 2024; Riecke et al., 2017). Prominent neurobiological accounts suggest that such attentional effects may be driven by high-level control regions in dorsal parts of the brain that modulate activity across sensory regions (e.g., Corbetta & Shulman, 2002; Yantis, 2008). Strikingly, recent data suggest that these control regions are not monolithic, with some dorsal subregions being recruited for visuospatial attention and other subregions exhibiting strong functional preferences for auditory tasks (e.g., Michalka et al., 2015; Noyce et al., 2017). While previous data suggest that voice and linguistic dimensions are largely analyzed by different neural systems (Hickok & Poeppel, 2007; Luthra, 2021; Maguinness et al., 2018), the present results indicate that differences in either dimension can modulate early attentional processing, likely due to extensive feedback circuitry in the brain.

The current work also raises several questions to be addressed by future work. For example, here we used relatively coarse semantic differences between streams (e.g., {*ba, da, ga*} vs {*1, 2, 3*}) and employed target syllables that do not have obvious semantic referents. It is an open question whether more subtle semantic differences (e.g., between animals and tools) would similarly support early filtering. Additionally, conditions were randomly intermixed across trials. This prevented participants from adapting to the challenge of ignoring a distractor that was similar in both voice and linguistic content. Future work might examine the extent to which listeners can adapt to the difficulty of ignoring highly similar distractor stimuli and how such adaptation might modulate the neural encoding of target stimuli. Finally, the present study focused on the mechanisms that support volitional, goal-directed control of attention. However, attention can also be involuntarily captured by salient stimuli, such as a cat meowing or a door slamming (e.g., Liang et al., 2022; Gaspelin & Luck, 2018; Theeuwes, 2023). Voice differences can provide a strong non-spatial cue to help listeners recover from the impact of unexpected interruptions and re-enage top-down attention (Liang et al., 2025). Thus, future work might examine how spatial, voice, and linguistic cues not only support top-down attention but also interact with bottom-up attention.

Overall, the present results provide insight into how listeners filter out irrelevant stimuli in cocktail party type environments. While voice or linguistic differences alone can facilitate spatial attention to a target stream, both features robustly influence the early neural encoding of target stimuli. Results highlight that when targets and distractors are similar, attention is less effective, leading to suppressed brain responses.

## Acknowledgements

This work was supported by a grant from the Office of Naval Research (ONR N00014-23-1-2065, PI: Shinn-Cunningham). SL was supported by NIH NRSA F32DC020625. GG was supported by a Summer Undergraduate Research Internship Experience in Acoustics from the Acoustical Society of America. We thank Iris Ta for assistance with data collection as well as Victoria Figarola and Benjamin Richardson for assistance with experiment programming and analyses.

## Author Contributions

Conceptualization: SL, WL, ALN, BGSC

Methodology: SL, WL, ALN, BGSC

Software: SL, HSK, WL

Formal analysis: SL, EP, MB, GG

Investigation: SL, EP, MB, GG

Resources: BGSC

Writing - Original Draft: SL, EP, MB, GG, HSK, WL, ALN, BGSC

Visualization: SL, ALN

Supervision: SL, ALN, BGSC

Project administration: SL, ALN, BGSC

Funding acquisition: BGSC, SL, GG

