## Supplemental Information for "Voice and linguistic features support early attentional filtering of irrelevant stimuli"

**Supplemental Behavioral Analyses**

In the main text, we analyze behavioral data from the target recall task considering only the primary fixed factors of Distractor Voice (same / different, deviation-coded with a [0.5, -0.5] contrast)) and Distractor Content (same / different, deviation-coded with a [0.5, -0.5] contrast). We obtained similar results in a more complex analysis that also considered possible interactions with Syllable Position (mean-centered) and Target Position (target leads / target lags behind distractor; deviation-coded with a [-0.5, 0.5] contrast). Results are summarized in Table S1.

**Table S1**

*Behavioral results evaluating target recall as a function of distractor voice, distractor content, syllable position, and target position*

| **Effect** | **b Estimate** | **Standard Error** | **z value** | **p value** | **Significance** |
|---|---|---|---|---|---|
| (Intercept) | 1.728 | 0.120 | 14.430 | < 0.001 | *** |
| Voice | -0.140 | 0.019 | -7.253 | < 0.001 | *** |
| Content | -0.558 | 0.019 | -28.827 | < 0.001 | *** |
| Syllable Position | -0.233 | 0.010 | -24.156 | < 0.001 | *** |
| Target Position | 0.005 | 0.019 | 0.283 | 0.777 | |
| Voice x Content | -0.540 | 0.039 | -13.946 | < 0.001 | *** |
| Voice x Syllable Position | -0.023 | 0.019 | -1.193 | 0.233 | |
| Content x Syllable Position | 0.068 | 0.019 | 3.544 | < 0.001 | *** |
| Voice x Target Position | 0.021 | 0.039 | 0.531 | 0.596 | |
| Content x Target Position | -0.372 | 0.039 | -9.606 | < 0.001 | *** |
| Syllable Position x Target Position | 0.167 | 0.019 | 8.668 | < 0.001 | *** |
| Voice x Content x Syllable Position | -0.056 | 0.039 | -1.449 | 0.147 | |
| Voice x Content x Target Position | -0.053 | 0.077 | -0.682 | 0.495 | |

| | | | | | |
|---|---|---|---|---|---|
| Voice x Syllable Position x Target Position | 0.018 | 0.039 | 0.459 | 0.646 | |
| Content x Syllable Position x Target Position | 0.383 | 0.039 | 9.907 | < 0.001 | *** |
| Voice x Content x Syllable Position x Target Position | 0.022 | 0.077 | 0.281 | 0.779 | |

*Estimates and standard error values are expressed on a log-odds scale. *** indicates p < 0.001, ** indicates p < 0.01, * indicates p < 0.05*

We found that task accuracy was reduced when the distractor was produced by the same voice as the target (main effect of Voice) or contained similar content (main effect of Content), and accuracy was generally worse for syllables that occurred later in the target stream (main effect of Syllable Position). Of primary theoretical interest, we also obtained a significant Voice x Content interaction: When listeners heard the same content in both streams, hearing the same voice in both streams led to an accuracy cost, but this was not the case when the content differed between streams.

In addition to this crucial interaction, we also observed a significant three-way interaction between Content, Syllable Position, and Target Position (as well as all two-way interactions between these factors), as visualized in Figure S1. Follow-up analyses indicate that listeners endured an accuracy cost if the trial began with a distractor that was linguistically similar to the target stream. We interpret these results as evidence that when a trial began with a distractor that was linguistically similar to the target, listeners' attention may have been captured by the distractor; the cost of reengaging attention manifests as relatively poorer detection of the first target stimulus on that trial.

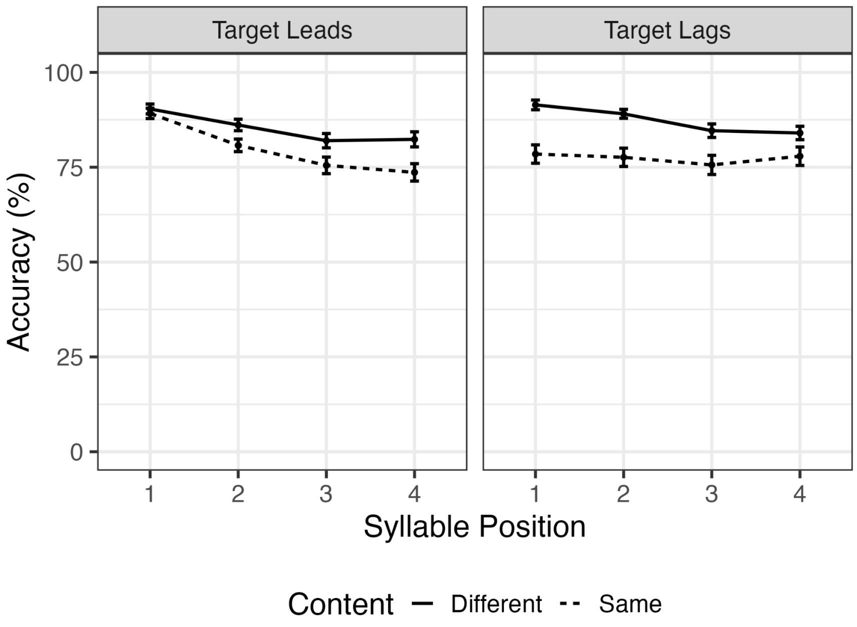


**Figure S1**. Percent accuracy in target recall (y-axis), as a function of syllable position (x-axis), target position (target leading / lagging behind distractor; left vs. right panel), and distractor content (solid vs dashed line). Follow-up analyses indicate that listeners endured an accuracy cost if the trial began with a distractor that was linguistically similar to the target stream.

**Supplemental ERP Analyses**

We analyzed the time course of ERP responses (separately for target and distractor ERPs) using cluster-based permutation testing implemented with the *clusterperm.lmer* function in the “permutes” library (Voeten, 2023). These analyses included fixed effects of Voice, Content, and their interaction, as well as random by-subject intercepts. The output is a set of time windows and associated cluster-mass statistics. To estimate permutation p values for the time windows that achieved significance, we ran follow-up regression analyses using the *perm.lmer* function that considered only the data from that time window, aggregated across time.

Event-related responses to the targets were significantly larger when the distractor was produced by the same voice as the target compared to by a different voice (main effect of Voice: 0 ms to 297 ms, $p < 0.001$), and amplitudes were larger when both target and distractor streams comprised syllables compard to when the distractor stream was composed of digits (main effect of Content:47 ms to 297 ms, $p < 0.001$). We also observed a significant interaction between these factors from 246 ms to 273 ms ($p = 0.003$). However, we interpret these results with caution. Because the target and distractor stimuli directly abut each other (i.e., there is a 0-ms ISI), by-condition differences in the time course of target ERP responses may be driven by responses to the preceding distractor. We therefore focus our interpretation on the analysis of peak-to-peak differences reported in the main text, since carryover effects of preceding distractors would be subtracted out in a peak-to-peak analysis.

Finally, although we were principally interested in effects on neural encoding of the target, we also examined event-related responses to distractor stimuli (Figure 3A, red panel). Visual inspection of the data suggests that the timing of ERP components differed across conditions, precluding a principled analysis of potential amplitude differences across conditions. Accordingly, we conducted a cluster-based permutation analysis to identify time windows in which neural responses to the distractor differed significantly across conditions. Results indicated that when distractors were linguistically similar to the target, they elicited strong, early neural responses (significant effect of content from 35 ms to 191 ms, $p < 0.001$); by contrast, distractors that were dissimilar to the target elicited strong responses only in a relatively late time window (significant effect of content from 203 ms to 297 ms, $p = 0.002$). The late portion of the ERP timecourse was also characterized by larger responses to distractors produced by a different voice relative to same-voice distractors (significant effect of voice from 230 ms to 297 ms, $p < 0.001$). Finally, we observed two portions of the time course where ERPs were influenced by both the voice and lingustic content (significant Voice x Content interaction from 172 ms to 242 ms, $p < 0.001$, and from 258 ms to 297 ms, $p < 0.001$). These results indicate that voice and linguistic features may influence attention at different time scales. However, because distractor stimuli differ acoustically across all conditions, the present design does not allow us to definitively ascertain whether these effects arise due to acoustic or attentional differences across conditions; accordingly, we consider these analyses to be exploratory in nature.
